# Engraftment and injury repair in regionally conditioned rat lung *in vivo* by lung progenitors derived from human pluripotent stem cells

**DOI:** 10.1101/2023.11.28.569060

**Authors:** Hsiao-Yun Liu, Camilla Predella, Ya-Wen Chen, Jing Wang, Mikael Pezet, Songjingyi Liang, Silvia Farè, John W. Murray, Anjali Saqi, Gordana Vunjak-Novakovic, Hans-Willem Snoeck, N. Valerio Dorrello

## Abstract

Although lung disease is a major cause of mortality, the mechanisms involved in human lung regeneration are unclear because of the lack of experimental models. Here we report a novel model where human pluripotent stem cell-derived expandable cell lines sharing features of airway secretory and basal cells engraft in the distal rat lung after conditioning by locoregional de-epithelialization followed by irradiation and immunosuppression. The engrafting cells, which we named distal lung epithelial progenitors (DLEPs), contributed to alveolar epithelial cells and generated ‘KRT5-pods’, structures involved in distal lung repair after severe injury, but only rarely to distal airways. Most strikingly, however, injury induced by the conditioning regimen was largely prevented by the engrafting DLEPs. The approach described here provides a model to study mechanisms involved in human lung regeneration, and potentially lays the foundation for the preclinical development of cell therapy to treat lung injury and disease.

## INTRODUCTION

Lung disease is a leading cause of morbidity and mortality. However, the lung has an extensive regenerative capacity that could be harnessed to develop disease-modifying treatments. The repair mechanisms remain unclear, especially in humans.

Mouse studies have shown multiple layers of often facultative progenitors in the distal lung.^1^ Although a small fraction of type 2 alveolar epithelial (AT2) cells^2–4^ can differentiate into type 1 (AT1) cells, likely through intermediates described as pre-AT1 transitional state (PATS)^5^ and ‘alveolar differentiation intermediate’ (ADI)^6^ cells, several other regenerative populations have been described in the distal lung. After partial pneumonectomy, AT1 cells can participate in alveolar regrowth.^7^ A ‘bronchioalveolar stem cell’ that co-expresses SCGB1A1 (a secretory cell marker) and SFTPC (an AT2 marker) may contribute to alveolar regeneration as well.^8,9^ Airway secretory cells, or subpopulations thereof,^10,11^ can convert to AT2 cells,^12,13^ although other studies showed no contribution of *Scgb1a1*-traced cells to alveolar repair.^14^ After severe injury, a population of p63^+^KRT5^+^ cells resembling airway basal cells (BCs) migrates distally as ‘KRT5-pods’ and participates in repair.^15,16^ These cells originate at least in part from a heterogeneous population of lineage-negative epithelial progenitors (LNEP),^17^ that also includes a population lineage-traced by the club cell marker, *Scgb1a1*, and expresses ITG4, CD200, and high levels of MHC class 2.^18^

Single cell transcriptomic studies have suggested that AT2-to-AT1 plasticity also exists in humans.^19–22^ Similar to mice, cells that share characteristics with airway BCs and secretory cells play a role in human lung regeneration. Kumar et al. cultured p63^+^KRT5^+^ ‘distal airway stem cells’ with alveolar potential *in vitro*,^15^ and accumulation of KRT5^+^ cells in the distal lung was observed in postmortem samples after severe injury.^23^ In contrast to mice, primates and ferrets have ‘terminal respiratory bronchioles’ (TRBs), small airways into which individual alveoli open, that terminate in alveolar ducts, where multiple alveoli coalesce. Cell populations expressing *SCGB3A2* (a secretory cell marker) and SFTPB (an AT2 marker) have been identified in TRBs. Based on organoid studies, bioinformatic trajectory analysis, and their higher abundance after lung injury, these TRB cell populations are believed to play a role in distal lung regeneration.^19–21^ Gaining functional insight into the role of regenerative populations after lung injury in the human lung has been challenging, since isolation and expansion of relevant cell populations from human lungs have not been accomplished yet, and functional studies *in vivo* require animal models engrafted with human cells. Nikolić et al. could achieve some engraftment in airways of bleomycin-treated immunodeficient *Nod.Scid.il2rg^-/-^* (NSG) mice with expanded putative human fetal lung distal tip cells, but they only followed up to 8 days and did not monitor differentiation.^24^ Fetal lung cells are additionally difficult to access. An approach to generate regenerative populations from human pluripotent stem cells (hPSCs) and to test their potential and fate *in vivo* would greatly advance this important area of research. Cells generated from hPSCs with the phenotype of fetal distal tip cells showed some engraftment potential in the airways of naphthalene-treated NSG mice.^25–27^ However, engraftment of human cells into the distal lung or promotion of lung repair by the engrafting human cells has never been definitively demonstrated.

Our study was designed to fill this gap by developing a xenogeneic rat model that allows engraftment and lung repair by expandable human cells with characteristics of both BCs and secretory cells generated from human pluripotent stem cell (hPSC)-derived lung organoids.

## RESULTS

### Generation of hPSC-derived lung progenitors

From lung organoids derived from human pluripotent stem cells (hPSCs), comprising both embryonic stem cells (ESCs) or induced pluripotent stem cells (iPSCs),^28–31^ we generated expandable cell lines by growing single cell suspensions in 2D on feeders **(Fig. 1a)**. Lines were expanded exponentially for at least 8 passages **(Fig. 1b)** and maintained normal karyotype **(Extended Data** Fig. 1a). Single cell RNA sequencing (scRNAseq) showed several clusters (differentially expressed genes shown in **Suppl. Table 1**, one of which were the feeder cells, which were discarded from further analysis) **(Fig. 1c)**. While it was challenging to assign cell types to each cluster, supervised analysis allowed subsetting into two major clusters based on expression of NOTCH ligands and receptors: BC-like and secretory-like receptors **(Fig. 1c-e)**. These were found in both ESC and iPSC-derived lines **(Extended Data Fig. 1b)**, and analysis was therefore integrated **(Fig. 1c**, right panel). The secretory-like cluster expressed *NOTCH3*, *UPK2*, *UPK1A*, *UPK1B*, *UPK3A*, *KRT4*, and *KRT13* with rare cells expressing the mature secretory marker, *SCGB1A1* **(Fig. 1d**, select feature plots shown in **Fig. 1e)**. These markers are expressed in secretory airway populations according to LungMap.^32^ *KRT4* and *KRT13* are also expressed in airway ‘hillocks’ in the mouse and are believed to an intermediate stage between BCs and secretory cells.^33^ In the mouse, *UPK3A* is expressed in rare variant club cells that reside near neuroendocrine bodies.^10,34^ *UPK3A^+^* variant secretory cells are remarkably increased in human neuroendocrine hyperplasia of infancy (NEHI).^11^ The secretory-like population also expressed *NOTCH2* (not shown) and *NOTCH3* **(Fig. 1d)** and showed higher expression of the NOTCH target gene, *HES4* **(Fig. 1d)** compared to the other clusters. These findings are consistent with the requirement for NOTCH signaling in the secretory lineage^35,36^ and with the observation that NOTCH signaling induces *UPK3A* in mouse airways.^10,34^ The second, larger population resembles airway BCs by virtue of expression of the BC markers, *TP63*, *KRT5*, *KRT17,* and *ITGB4* **(Fig. 1d)**. This population expressed the NOTCH ligands, JAG2 and DLK2 **(Fig. 1d)**, suggesting mutual regulation of both fates. Expression of KRT5, KRT17, and p63 were confirmed by immunofluorescence (IF) **(Fig. 1f)**, whereas a population of MUC1^+^ cells was detected by flow cytometry **(Fig. 1g)**. Consistent with an airway BC-like cell phenotype, the cells generated airway cells (ciliated, goblet and club cells) when cultured in air-liquid interface conditions, albeit with cell line-dependent kinetics **(Extended Data Fig. 1c)**.

**Figure 1.**
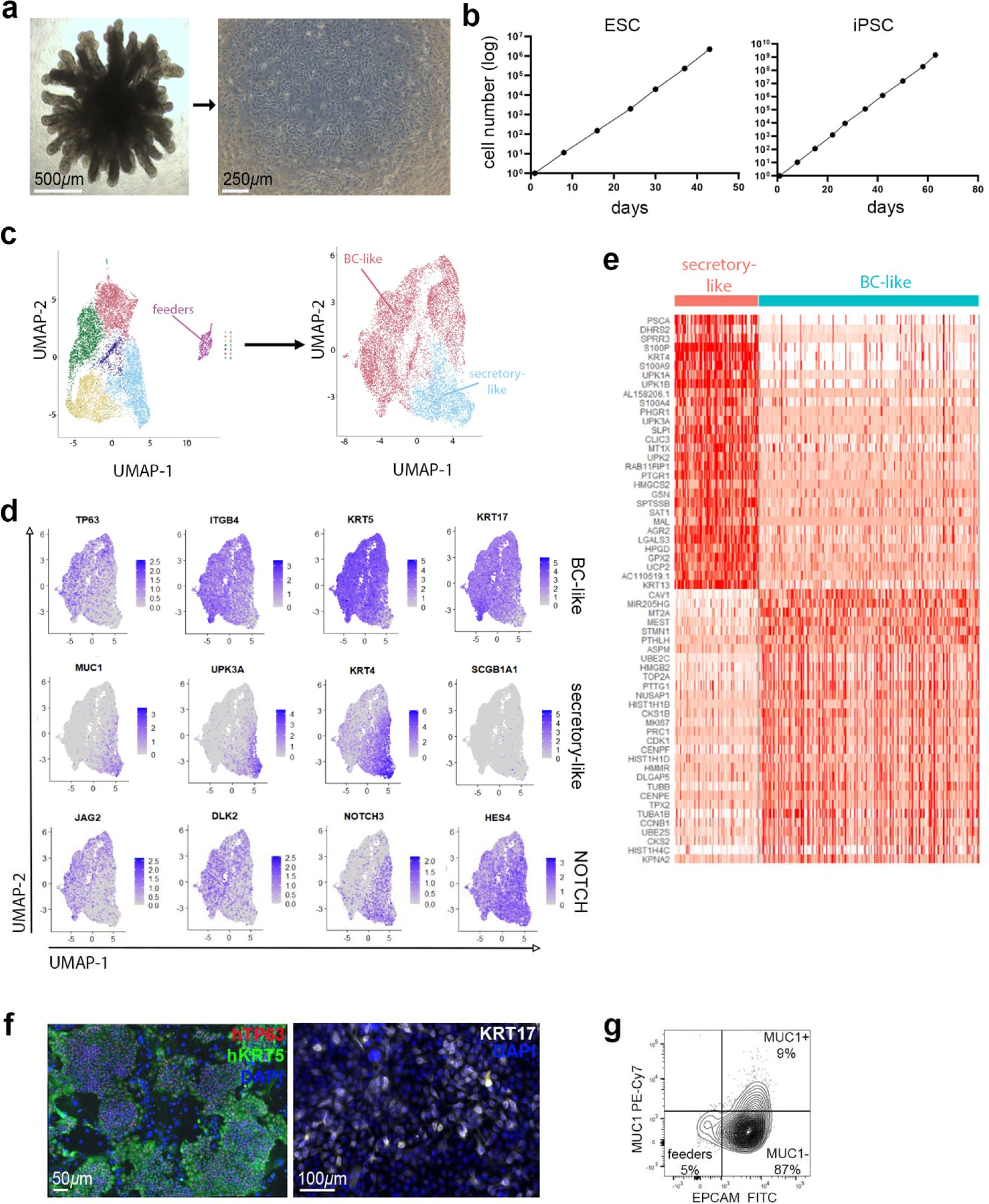
Characterization of lung progenitor lines. **a.** Generation of DLEPs from 3D organoids**. b.** Expansion of DLEPs generated from an ESC and an iPSC line. **c.** UMAP of scRNAseq data including feeder cells (left), and integrated UMAP for ESC and iPSC-derived DLEPs after subsetting based on NOTCH ligand/receptor expression (right). **d.** Feature plots for genes associated with BCs, secretory cells, and NOTCH signaling. **e.** Top differentially expressed genes in both subsets. **f.** IF for hTP63, hKRT5, and KRT17 in ESC-derived DLEPs. **g.** Flow cytometric analysis of MUC1 expression in DLEPs.

### Development of a lung injury model in rats

In immunodeficient NSG mice treated with either naphthalene or bleomycin, with or without irradiation to block endogenous repair, we could not achieve reproducible engraftment with aforementioned cells, although some human cells were occasionally observed. We therefore developed a model based on locoregional de-epithelialization, which is not technically feasible in mice, in a larger animal, the rat.

We used a mild detergent solution (see Online Methods), previously shown to locally remove lung epithelium in *ex vivo* ventilated rat lungs.^37^ An engineered cannula with a proximal plastic plug was lodged in the lower left lung of an anesthetized, ventilated, and monitored rat using a flexible bronchoscope thus creating a temporary seal in the trachea to facilitate removal of de-epithelialization solution **(Extended Data Fig. 2a,b)**. De-epithelialization solution was administered followed by saline wash **(Extended Data Fig. 2c)**. Blue food dye was added in the initial experiments to macroscopically identify the treated area **(Extended Data Fig. 2d)**. Heart rate and oxygen saturation decreased during the procedure but returned within normal range after de-epithelialization **(Extended Data Fig. 2e)**. Hematological, hepatic, metabolic, and renal values were within the normal range 48 hours post de-epithelialization **(Extended Data Fig. 2f)**. None of the animals showed any sign of respiratory distress at 48 hours or later time points, although we observed approximately 5% animal mortality due to complications related to anesthesia and/or intubation.

### Validation of *in vivo* de-epithelialization

Compared to the untreated right lung within the same animal, there was a considerable loss of AT1 (^+^) and AT2 (proSPC^+^) cells in the treated left lung region 48hrs post de-epithelialization as indicated by IF, whereas expression of endothelial marker, CD31, was preserved, indicating mostly selective removal of the epithelium **(Fig. 2a)**. Hematoxylin and eosin (H&E) staining showed patency of the alveoli in the targeted region **(Fig. 2a)**. Regional de-epithelialization was further confirmed by immunohistochemistry (IHC) showing lack of staining for pan-keratin (Pank), but maintenance of CD31 staining in the targeted region **(Fig 2b)**. The lung injury score (LIS), assessed according to ATS guidelines^38^ was significantly elevated in the lower left lung (treated) compared to the right lung (untreated) within the same animal **(Fig. 2c,d, Extended Data 3a,b)**. Neutrophils in alveolar walls and interstitial space, hyaline membranes, and thickening of septa were detected across the treated left lower lobe **(Fig. 2c,d** and **Extended Data Fig. 3a,b)**. We also observed a lower amount of proteinaceous debris in the de-epithelialized lungs compared to untreated lungs **(Fig. 2d)**, likely explained by the abundant presence of hyaline membranes in de-epithelialized lungs obscuring the underlying proteinaceous debris.

**Figure 2.**
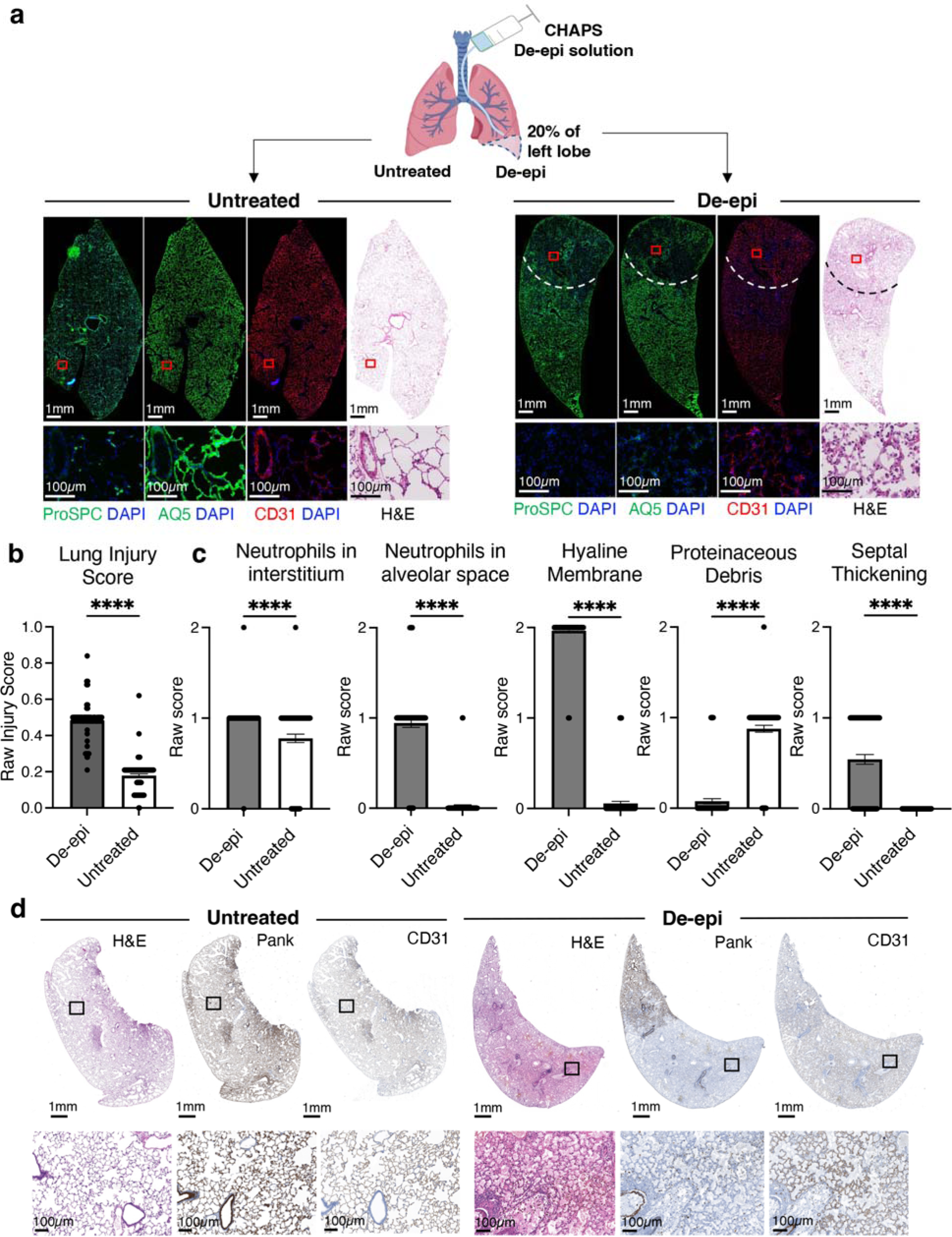
Regional de-epithelialization characterization. **a.** Schematic representation of the targeted region and IF staining for ProSPC (AT2), AQ5 (AT1), and CD31 (endothelial cells). De-epithelialized area outlined by the dotted white lines. Representative red squares are shown at higher magnification (lower panels). **b.** H&E, Pankeratin (Pank) and CD31 immunohistochemistry. Representative black squares are shown at higher magnification (lower panels). **c.** Lung injury score (LIS) of de-epithelialized region compared to the contralateral, untreated lung. **d.** Scores for individual features of the LIS. In both c and d, data are reported as mean±SEM and analyzed by paired t-test [n=3 animals per condition, 60 ROI (30 from left lung, 30 from right lung) per animal; ****, p<0.0001].

EdU labeling 48hrs after de-epithelialization showed extensive proliferation in the epithelium of airways and the alveoli in the targeted region as well as in the surrounding areas of the lower left lung **(Fig. 3a)**, indicating a regional regenerative response of the lung to the local injury **(Fig. 3a)**. After 5 days, partial recovery of alveolar epithelium occurred in the treated regions as evidenced by increased expression of and proSPC **(Fig. 3b)**. At day 10, the alveolar epithelium appeared fully reconstituted **(Fig. 3c)**. Assessment of the LIS **(Fig. 3d,e** and **Extended Data Fig. 3b)** and IHC confirmed these findings **(Fig. 3f,g)**.

**Figure 3.**
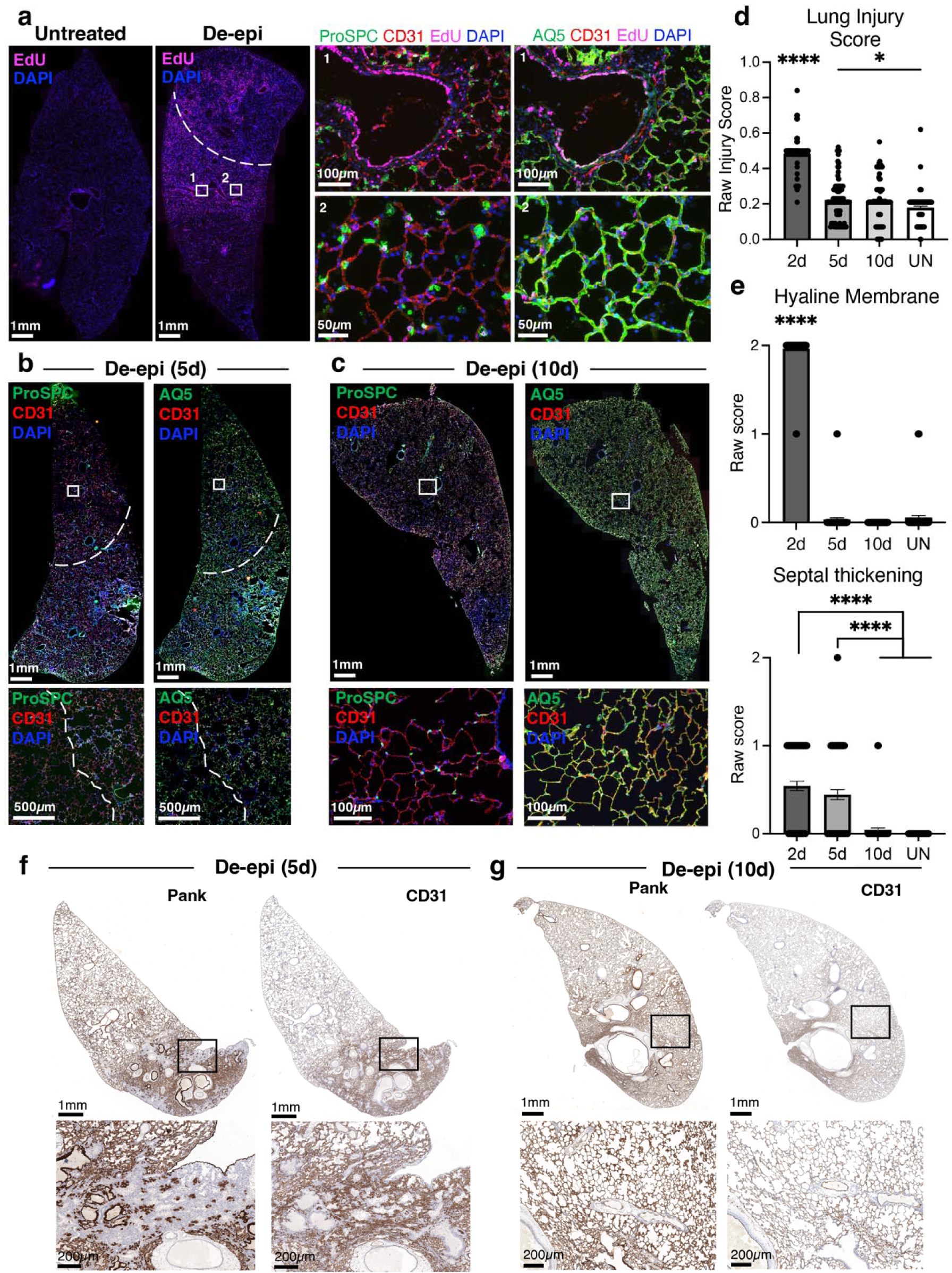
Endogenous re-epithelialization. **a.** EdU incorporation in the de-epithelialized region and in the contralateral lung. **b-c.** Immunofluorescent staining for epithelial cells (AQ5, ProSPC), and endothelial cells (CD31) in treated regions 5 (**b**) and 10 (**c**) days post de-epithelialization. **d.** LIS of de-epithelialized region evaluated 2-, 5-, and 10-days post-treatment and compared to untreated animals. Mean±SEM, one-way ANOVA, n=3 animals per condition, 60 ROI (30 from left lung, 30 from right lung) per animal; *, p=0.0287; ****, p<0.0001**. e.** Scores for the most pronouncedly different individual features of the LIS. Mean±SEM, one-way ANOVA, n=3 animals per condition, 60 ROI (30 from left lung, 30 from right lung) per animal; ****, p<0.0001. **f-g.** Pankeratin (Pank) and CD31 immunohistochemistry 5 days (**f**) and 10 days (**g**) after de-epithelialization. Representative black squares in lower panels.

**Engraftment of hPSC-derived lung progenitors**.

Next, we applied de-epithelialization, followed by an immune suppression regimen (see Online Methods). 48hrs after de-epithelialization, we administered 10^7^ hPSC-derived lung progenitors, or saline, intrabronchially targeting the lower region of left lung **(Fig. 4a)**. The observation time was limited to 10 days since daily subcutaneous injections of tacrolimus, required for appropriate immunosuppression, became more challenging after that because of induration of the injection site. After 10 days, patches of human cells organized as alveolar ‘nets’ **(Fig. 4b)**, were found scattered throughout the lower left lobe as judged by staining for human mitochondria (hMit) **(Fig. 4, Extended Data Fig. 4** and **Extended Data Fig. 5)**. Co-staining for proSPC, SPC, or RAGE indicated AT1 and AT2 development from the engrafted cells **(Fig. 4, Extended Data Fig. 4** and **Extended Data Fig. 5)**. No contribution to large or small airways was observed, however. Because of their capacity to engraft in the alveolar epithelium, we refer to these cells as ‘distal lung epithelial progenitors’ (DLEPs).

**Figure 4.**
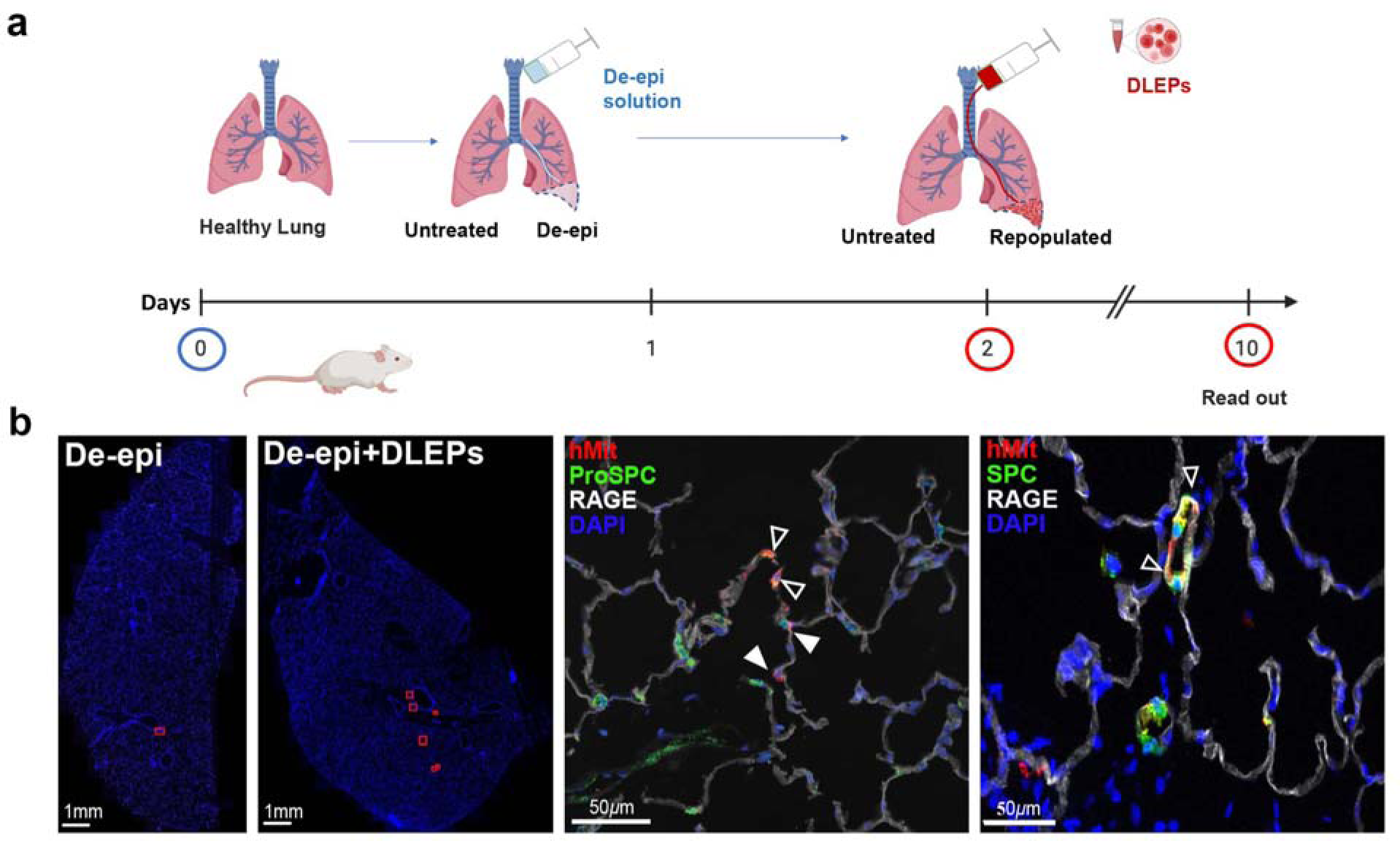
Engraftment of DLEPs in de-epithelialized rat lungs *in vivo*. **a.** Schematic representation of experimental approach**. b.** Annotations (red squares) indicating the location of human cell based on hMit staining, using de-epithelialized, non-engrafted lung as background control for IF. **c.** Representative areas of engrafted DLEPs detected by hMit, SPC, ProSPC, and RAGE (Yellow arrowheads: human cells co-expressing hMit with SPC or ProSPC. White arrowheads: human cells expressing hMit and RAGE). Single channels and a representative higher magnification area of engrafted cells are highlighted in the white squares.

To improve engraftment, we blocked endogenous lung repair with 12Gy local irradiation of the lung field 24 hours after de-epithelialization **(Extended Data Fig, 6a,b)**, which abolished EdU incorporation after injury **(Extended Data Fig.6c)**. Transplantation of DLEPs at day 2 **(Fig. 5a)** resulted in extensive areas containing human alveolar epithelium detected by staining for hMit, proSPC, SPC, and RAGE **(Fig. 5b, Extended Data Fig. 7a,b, Extended Data Fig. 8** and **Extended Data Fig. 9a)**. RNAscope^39^ for human β2-microglobulin (hB2M) confirmed patchy locoregional replacement of rat alveoli by human cells that corresponded to the staining for hMit **(Fig. 5c)**. We occasionally observed engraftment in airways, as identified by staining for hKRT5, hTP63 and hMit **(Extended Data Fig. 9b)**. We did not, however, find markers of mature cells at the 10-day time point. In addition, we observed dense aggregates of human cells, as judged by staining for hMit and RNAscope for hB2M, in localized areas that appeared histologically more severely damaged. These structures stained with human-specific anti-KRT5 anti-TP63 antibodies **(Fig. 5c, right panels** and **Extended Data Fig. 9c)**, and are therefore highly reminiscent of the KRT5-pods identified in the distal lungs of mice after severe influenza.^15–17^

**Figure 5.**
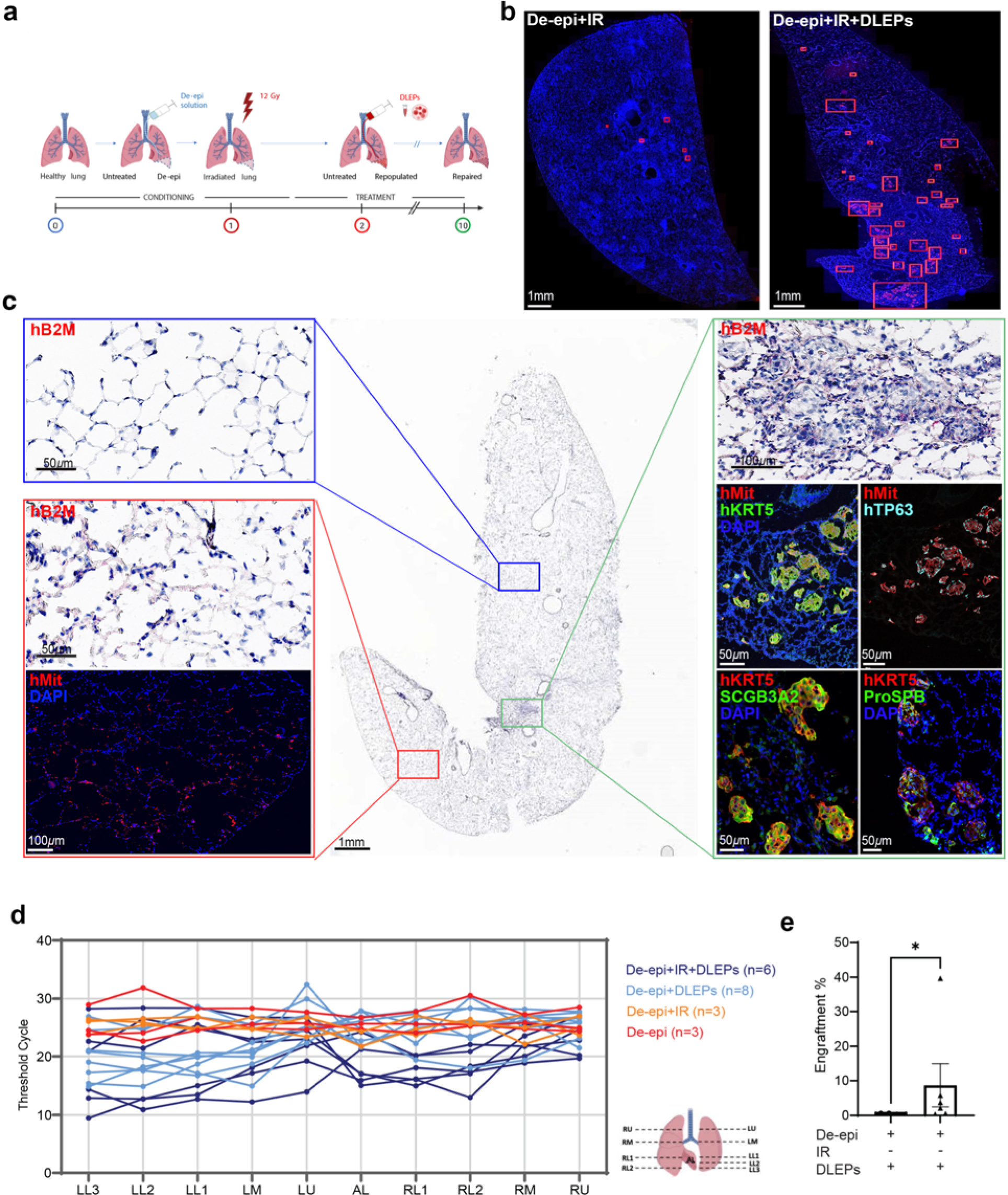
Engraftment of DLEPs after de-epithelialization and irradiation. **a.** Schematic representation of experimental approach. **b.** Annotations (red squares) of the location of human cells based on hMit staining, using de-epithelialized, non-engrafted lung as background control. **c.** RNA *in situ* hybridization (RNAscope) for human b2-microglobulin (hB2M), and IF for hMit, hKRT5, hTP63, ProSPB, ProSPC, RAGE and SCGB3A2 in representative engrafted area from left lung. (Yellow arrowheads: human cells co-expressing hMit with ProSPC). Higher magnification areas of engrafted cells are highlighted in the white squares. The left upper panel shows a region without human cells as a negative control for RNAscope. **d.** Human gDNA content in sections of lung regions as represented in the schematic on the right. Aggregated data from equal numbers of experiments with ESC- and iPSC-derived DLEPs. **e.** Percent engraftment in the most highly engrafted section of panel **d** for each experiment. Mean±SEM, unpaired Student’s t-test (n=6 per condition; *, p=0.02).

Furthermore, the human engrafted ‘KRT5-pods’ expressed SCGB3A2, proSPC, and proSPB **(Fig. 5c, lower right panels** and **Extended Data Fig. 9c)**, markers co-expressed in recently identified putative distal lung progenitors in human lungs. ^20,21^

To estimate the number of engrafted human cells, genomic DNA (gDNA) was extracted from ten 200µm thick sections from right and left lungs according to the schematic in **Fig. 5d** and subjected to qPCR for human AluYb8 **(Extended Data Fig. 10a)**.^40^ Using a standard curve of varying amounts of human cells alone or mixed with rat lung cells **(Extended Data Fig. 10b)**, we estimated engraftment of human cells within the section with the highest engraftment in each individual animal **(Fig. 5d)**. In de-epithelialized/transplanted region, we found ∼0.6% human cells per section **(Fig. 5e)**. With irradiation, however, average engraftment was ∼10%. **Fig. 5d** also shows that in occasional rats, the right lung was injured and engrafted as well, most likely due to spillover or to anatomical variation in the airways of these outbred rats, which could lead inadvertent targeting of the right lung.

We conclude that DLEPs can engraft in rat lungs after appropriate conditioning and immunosuppression following three patterns of engraftment: KRT5-pods in severely damage areas, integrated replacement of alveolar epithelium, and, occasionally, engraftment of airways.

### Injury repair by DLEPs

Low-magnification images of HE-stained sections showed that the combination of de-epithelialization and irradiation induced severe damage of lung epithelium **(Fig. 6a,b)**. The lower left lobes of rats engrafted with cells, however, showed a remarkable attenuation of lung damage, although some localized injury was still observed **(Fig. 6a,b)**. To quantify injury and the effect DLEPs on injury repair, we assessed the LIS. Rat lungs receiving either ESC or iPSC-derived lines showed significant reduction in LIS that, importantly, quantitatively approached that of uninjured lungs **(Fig. 6b** and **Extended Data Fig. 10c)**. The most strikingly elevated feature of the LIS in the conditioned lower left lobe was septal thickening **(Extended Data Fig. 10c)**. This feature, in particular, was reduced to near uninjured animals’ levels in all transplanted recipients **(Extended Data Fig. 10c)**. These data indicate that DLEP engraftment promotes repair, and therefore unequivocally showing functionality.

**Figure 6.**
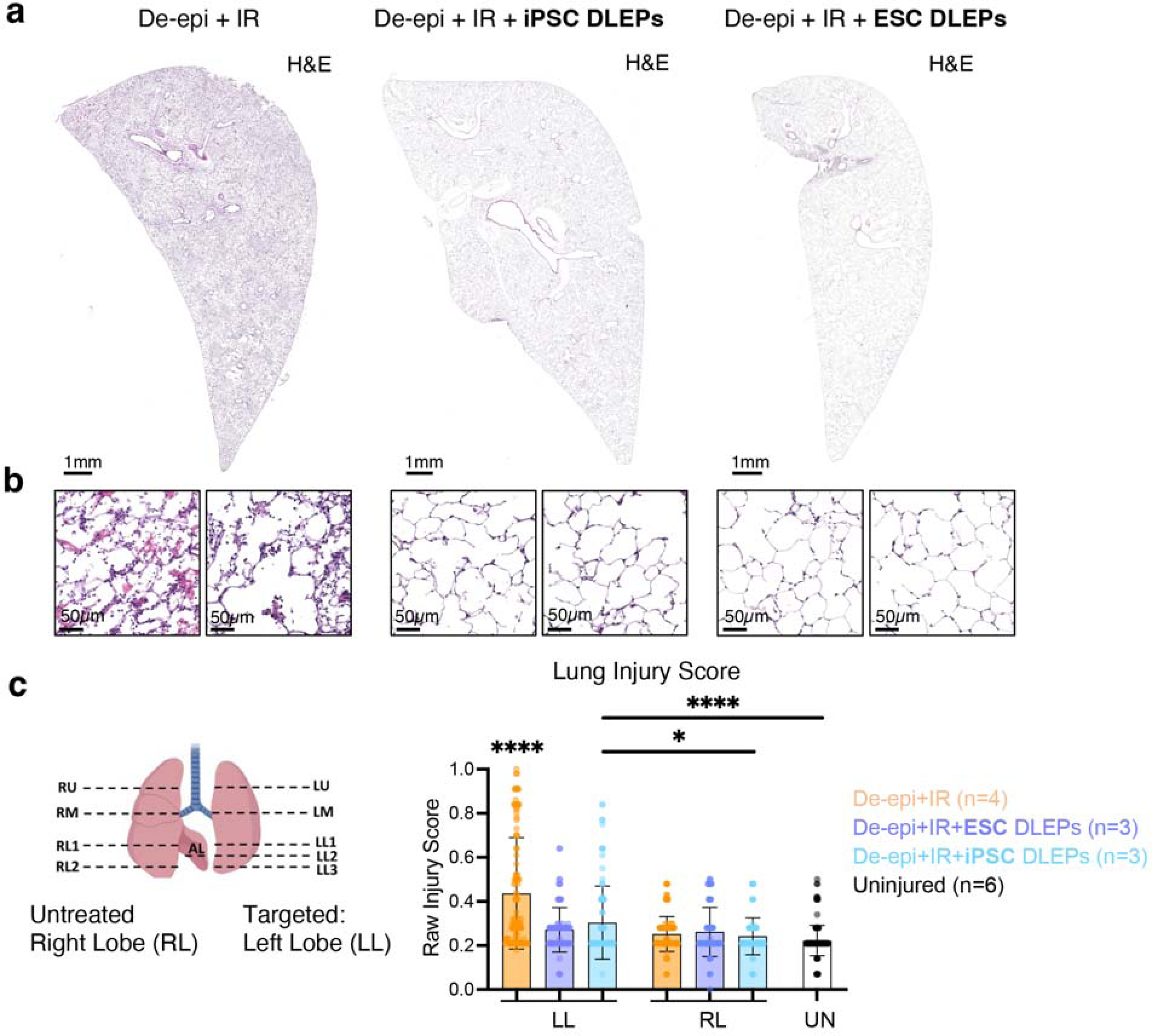
Repair of lung injury. **a.** H&E staining of injured (De-epi + IR) and engrafted (De-Epi + IR + DLEPs) lower left lobes. **b.** Representative higher magnification images. **c.** Lung injury scores (IR: irradiation, UN: normal control lung, 800 blindly evaluated fields in total, 30 ROI per tile scan, one-way ANOVA; *, p=0.0383; ****, p<0.0001).

## DISCUSSION

We report here that hPSC-derived cells engraft and integrate in the distal lung of rats conditioned with regional de-epithelialization followed by irradiation and promote repair of lung damage inflicted by the conditioning regimen, thus providing, to the best of our knowledge, the first strategy for the generation of distal lung progenitors from hPSCs that engraft into rodent lung and the first model to study the role of such cells in lung regeneration and repair.

Although we use, by necessity, a xenotransplant model, where it can be assumed that the actual engraftment potential of the cells was underestimated despite immunosuppression, our findings that a cell population sharing expression signatures with airway BCs or secretory cells engrafted in the distal lung and promoted repair supports recent reports describing subsets of cells with BC-like and secretory-like phenotypes participating in distal lung regeneration in mice and, by inference, in humans.^5,6,12,16–18,20,21^

Interestingly, DLEPs generated structures similar to KRT5-pods^15–17^ only after irradiation, which combined with de-epithelialization induced severe and lasting injury. This did not occur in rats that were only treated with de-epithelialization, which rapidly recover spontaneously. This finding is consistent with the appearance of KRT5-pods after severe lung injury in mice.^15–18^

Expansion of KRT5^+^ cells has also been demonstrated after severe lung injury in humans, although these data were only obtained from post-mortem specimens.^23^ These cells co-expressed SFTPC,^23^ consistent with the expression of SFTPC, in addition to SCGB3A2 and SFTPB in engrafted KRT5^+^ cells in our model. Conditional deletion of KRT5^+^ cells after injury in the mouse led to persistent damage after injury and reduced formation of alveolar ‘nets’ consisting of AT1 cells, indicating their essential role in repair.^16^ We observed formation of such donor-derived alveolar ‘nets’ in our model as well. Although the quantitative contribution of KRT5^+^ cells to the alveolar epithelium is unclear, KRT5^+^ cells arising after severe lung injury may also fulfill a supportive role, either structural or mediated by paracrine factors, in lung repair.^1,17^ Consistent with this idea, the extent of injury repair by engrafted DLEPs after conditioning with de-epithelialization/irradiation was much more extensive than the actual alveolar engraftment. Our data therefore suggest that DLEPs generated from hPSCs are potentially the precursors of the human equivalent of regenerative KRT5^+^ cells in mice and provide a unique *in vivo* model to further investigate the biology and fate of these cells.

Airway repopulation was rare and only observed in two recipients, although DLEPs have airway potential *in vitro*, as shown in ALI studies. It is possible that physical forces caused by spontaneous breathing and mechanical ventilation drive both the detergent solution and the cells to the distal lung, such that injury and cell attachment occur predominantly in the alveoli.

Our observations may also establish the basis for a pre-clinical model for cell therapy for lung diseases, in particular those characterized by acute injury of distal lung, such as in acute respiratory distress syndrome, or by an aberrant response to chronic injury, such as in idiopathic pulmonary fibrosis. Implementation of cell therapy for lung disease requires both appropriate engrafting cells that can be expanded at clinical scale and a safe strategy to condition the recipient lungs. The DLEPs we generated are one potential candidate population. AT2 cells may also constitute an interesting cell type, as these are facultative alveolar stem cells. Generation of AT2 cells from hPSCs requires the use of reporters or sequential sorting for surface markers, however, and the cell population obtained contains significant numbers of gastric cells as well as cells that do not express the mature AT2 markers, *SFTPB* and *SFTPC*, by scRNAseq.^41^ Engraftment or injury repair with such cells has not been reported. Interestingly, Vaughan et al. showed that mouse AT2 cells cultured and expanded as organoids gave rise to KRT5^+^SCGB3A2^+^ cells after transplantation into influenza-conditioned mouse lung without providing functional benefit.^42^ In contrast, fresh, primary murine AT2 cells did regenerate AT1 and AT2 cells, suggesting that extended culture and expansion of AT2 cells affected their regenerative potential.^42^

Mouse models of lung injury, including bleomycin, toxic gases, and naphthalene^43^ cannot be used clinically because of unacceptable toxicity. Two groups pioneered *ex vivo* lung decellularization followed by seeding primary epithelial cells into the airways and endothelial cells into the vascular compartment. However, these bioengineered lungs failed only a few hours after transplantation due to alveolar edema and thrombosis.^44,45^ Sequential regional, mild detergent-mediated de-epithelialization, as we demonstrated here, might provide a better avenue to develop cell therapy for lung injury and disease.

We propose that lung progenitors derived from human pluripotent stem cells as engrafting cells and the xenogeneic rat model with local conditioning described here represent a promising avenue to gain insight into human lung repair and to potentially to develop novel approaches to treat lung injury and distal lung disease.

## METHODS

### Animals, anesthesia, and euthanasia

Sprague-Dawley (SD) rats weighing 230-250 and 180-220 g were used in the experiments conducted to optimize and characterize the regional lung de-epithelialization and irradiation (n=36), and the transplant studies (n=14), respectively. All animal work was approved by the Columbia University Institutional Animal Care and Use Committee (IACUC), complying with the National Research Council Guide for the Care and Use of Laboratory Animals (eighth edition). Rats, both males and females, were anesthetized with isoflurane vapor (3-5%) and an intraperitoneal (IP) injection of ketamine (VEDCO, Saint Joseph, MO) and xylazine (Covetrus, Portland, ME). The animals were then endotracheally intubated with a modified cannula (JorVet, Loveland, CO) **(Extended Data4a,b)**. Throughout the whole experiment, animals, maintained on 1% isoflurane, were spontaneously breathing through the lung (right) not intubated and mechanically ventilated through the intubated lung (left). The left lung was ventilated using synchronized intermitted mandatory ventilation (SIMV) on volume control modality. Euthanasia was performed with Isoflurane (Covetrus) (5%, 15min) inhalation, followed by bilateral thoracotomy. Following a transverse incision of the aorta to facilitate blood release from left ventricle, the main pulmonary artery was perfused by phosphate-buffered saline (PBS, Sigma-Aldrich, St. Louis, MO) via the right ventricle with static pressure that was maintained at 13 cm H_2_O above the heart until lungs, perfused, were blanched. OCT compound (Sakura Finetek, Torrance, CA) was diluted in PBS (8 OCT:2 PBS in volume) and injected into the trachea. A total of ten tissue samples were taken from each animal and embedded in the undiluted OCT compound for frozen sections.

### Monitoring and blood analysis

Heart rate, oxygen saturation, and rectal temperature were monitored during each experiment using a pulse oximeter (8500, Nonin Medical, Plymouth, MN) and a digital thermometer, respectively. Tail vein blood was collected one day before and 48 hours after DE experiments and delivered to Columbia University Comparative Pathology Laboratory (CPL) for pathologic analysis. In the engraftment experiments, the plasma level of tacrolimus was measured in blood samples collected from experimental and control rats the day of cells delivery and at 7 days post-transplant. Plasma was processed with the Abbott specific kit for tacrolimus level, according to the kit manufacturer’s protocol (Abbott, Abbott Park, IL) and read with the Abbott Architect machine.

### Lung irradiation

One day after de-epithelialization, rats were anesthetized with Isoflurane (1-3%) and placed on the animal holder of the Small Animal Radiation Research Platform (SARRP). After the total body computerized tomography (CT) scan, the lungs area was contoured and selected for irradiation with a build-in software of the platform. Each rat was irradiated using two beams, one from top and one from bottom respect to the animal.

### hPSC maintenance

RUES 2 (Rockfeller University Embryonic Stem Cell Line 2, passage 20-27), Sendai Virus-induced human dermal fibroblast iPSC lines (from healthy fibroblasts, purchased from Mount Sinai Stem Cell Core Facility, passage 17-23) were cultured on mouse embryonic fibroblasts (GlobalStem, Gaithersbug, MD) plated at 17,000-20,000 cells/cm^2^. hPSC maintenance media consisted of DMEM/F12 (Croning, Teterboro, NJ), 20% Knockout Serum Replacement (Gibco), 0.1 mM β-mercaptoethanol (Sigma-Aldrich), 1% GlutaMax (Gibco), 1% non-essential amino acids (Gibco), 0.2% primocin (InvivoGen, San Diego, CA) and 20 ng/ml FGF-2 (R&D Systems, Minneapolis, MN). Media were changed daily, and cells were passaged every 4 to 5 days using Accutase/EDTA (Innovative Cell Technologies, San Diego, CA) and plated at 10,000-12,000 cells/cm^2^ density. Cells were maintained in an undifferentiated state in a humidified 5% CO_2_ atmosphere at 37°C. The cells were tested for Mycoplasma contamination by PCR every 6 months. Karyotype was performed every 6 months.

### Generation of hPSC-derived lung organoids and DLEPs

The hPSC-derived human lung organoids were generated as described.^31^ For DLEP generation, Matrigel-embedded lung organoids were released from Matrigel and grown in conditions conducive to growth of airway progenitors. DLEPs were passaged every week and could be kept more than 20 passages with normal karyotype.

### Cell transplantation

For the transplantation studies, rats were started on a daily triple drug immunosuppression regiment consisting of with Tacrolimus (Prograf from Astellas, Northbrook, IL), Mycophenolate (Accord, Durham, NC), and Methylprednisolone (Sagent, Schaumburg, IL) 24 hours after de-epithelialization and 24 hours prior to administration of the cells. Rats were anesthetized and intubated with the same modified cannula used for the regional de-epithelialization (JorVet) targeting the lower left lobe with of a flexible bronchoscope (1800Endoscope, Bradenton, Florida). 10^7^ cells (DLEPs resuspended in their culture media) were delivered through the cannula. Rats were weaned from anesthesia, returned to their cages, and euthanized as described 10 days from the de-epithelialization.

### Flow Cytometry analysis

Single-cell suspensions of DLEPs were obtained dissociating with 0.05% Trypsin/EDTA in normoxic incubator for 10-12 minutes with occasional pipetting with P1000. Dissociated cells were neutralized with wash media, then centrifuged at 400 g for 4 minutes. Cells were stained with conjugated antibodies (see **Table S2** for list of antibodies used) for 20 minutes in FACS buffer (1% BSA in PBS) at 4C and then resuspended in FACS buffer with 1:1000 DAPI for live/dead cells quantification. FACS was performed on BD® LSR II Flow Cytometer (BD, Franklin Lakes, NJ) and data analyzed with FlowJo 10.

### Air-liquid interphase culture

10^6^ DLEPs were seeded onto 24-well trans-well insert and kept with DLEP media on the upper and lower chamber overnight. Cells were further cultured with PneumaCult™-Ex Plus Medium (Stem Cell Technologies) on the upper and lower chamber 48 hours, after which ALI was induced and cells maintained in PneumaCult™-ALI media (Stem Cell Technologies).

### scRNAseq analysis

Single-cell gene expression profiles for ESC-derived and iPSC-derived DLEPs cells were generated with 10x Genomics Chromium Single-Cell 3′ RNAseq platform. The Columbia University Genomic core performed raw data processing with 10x Genomics Cell Ranger pipeline. The reads were mapped to a human reference genomes (GRCh38). Counts for each sample were analyzed with Seurat, separately. Cells with a unique molecular identifier (UMI) lower than 500 counts were filtered. Cells with high level of mitochondrial reads (>20% of counts) were removed. The counts were normalized, scaled and analyzed for principal component analysis (PCA) with default methods. The principal components (PC) were used to generate the uniform manifold approximation and projection (UMAP), find neighbouring cells and identify cell clusters using default Seurat parameters. During clustering the resolution was adjusted to 0.1. To estimate the level of contamination by feeder cells, known cell type specific marker genes (ZEB2, FGF7, ZEB1) were used and identified as expressed by a separate cluster of cells in both samples. The feeder-associated cluster was subsetted out and integrated analysis implemented in Seurat was repeated with identical parameters. Differentially expressed genes among clusters and sample types were identified with the FindMarkers function in Seurat, with a log fold-change threshold of 0.25 and statistical cutoff of adjusted P value (>0.5). Clusters were reordered based on top differentially gene expression similarities.

### EdU (5-ethynyl-2’-deoxyuridine) incorporation

50 mg/kg EdU (Click Chemistry Tools, Scottsdale, AZ) was administered intraperitoneally 3 hours before euthanasia to label proliferating cells. EdU incorporation was visualized using the Click-iT Plus EdU Alexa Fluor 555 Imaging kit (ThermoFisher, Fair Lawn, NJ), according to the manufacturer’s instructions.

### Histology and immunofluorescence (IF) staining

Frozen sections (5µm thick) and H&E stainings from each experiment were prepared by the Molecular Pathology Shared Resource (MPSR) facility at Herbert Irving Comprehensive Cancer Center (HICCC),Columbia University, New York. Upon returning to the room temperature, the sections were submerged in 4% paraformaldehyde for 10 minutes, washed with PBS for 5 minutes, and permeabilized in 0.25% Triton X-100/PBS solution for 20 min. After one hour of blocking in 10% donkey serum/PBS solution, EdU staining was performed in the experiments where *in vivo* labeling was performed. Subsequently, primary antibodies were added to the sections diluted in 5% donkey serum/PBS solution. The slides were then incubated overnight in the dark at 4°C. The next day, following three 10-minute washes in 0.025% Triton X (ThermoFisher) −100/PBS solution, the sections were treated with secondary antibodies (in 5% donkey serum/PBS) for a 1hr in the dark. Slides were again washed three times for 10mins in 0.025% Triton X-100/PBS solution. DAPI staining (1:1000 in PBS) was performed for 10mins and the slides washes for 5mins in PBS before being mounted and sealed with nail polish. A list of primary and secondary antibodies utilized is available in **Table S2**. IF images were taken and processed using a Leica DMi8 system and with the Leica Stellaris for confocal imaging.

### RNAscope

RNAscope stainings were performed according to the manufacturer’s instructions (ACD - a Bio-Techne brand, Newark, CA) using the following probe and reagents: Beta-microglobulin (B2M)–C2 (Cat No. 1211661-C2) with RNAscope 2.5 HD Duplex Detection Ki (Chromogenic). Briefly, tissue was freshly harvested and embedded into OCT and further stored at the −80. Tissues were sectioned in 5µm and fixed with ice cold 4% paraformaldehyde at 4°C for 15 min. Slides were dehydrated with ethanol and air-dried completely. A hydrophobic barrier was drawn around the tissue with ImmEdge Pen (Vector Labs, Newark, CA). Endogenous peroxidase activity was blocked with hydrogen peroxide for 10 min at room temperature (RT). Slides were treated with RNAscope Protease IV for 15-30 min at RT and then processed to run the RNAscope assay. We hybridized the probes, applied RNAscope signal amplifiers and labeled probes according to the manufacturer’s instructions. The images were taken with Leica AT2 bright field whole slide scanning system with 40x magnification.

### Genomic qPCR

Genomic DNA was extracted from 200-250µm-thick cryopreserved rat lung tissue using the Zymo Quick-DNA MicroPrep Kit (Zymo, Irvine, CA), according to the manufacturer’s instructions. Tissues samples were washed with water to remove residual OCT before lysing with Zymo Genomic Lysis Buffer. DNA concentration was assessed by absorbance in a spectrophotometer (Thermo Scientific NanoDrop2000c). Human and Rat DNA were verified by qPCR using human AluYb8 primers (*Alu,* Fw 5’-CGAGGCGGGTGGATCATGAGGT-3’, Rv 5’-TCTGTCGCCCAAGGCCGGAGCT-3’) and rat *aryl hydrocarbon receptor* gene (*Ahr* Fw 5’-TGCCCAGCAACAGCCTGTGAG-3’, Rv 5’-AACTGGCGAACATGCCATTGA-3’), as described by Prigent *et al.*^40^ Serially diluted human genomic DNA (0.01pg-10ng) was mixed with 100ng of rat genomic DNA to determine a standard curve at each set of qPCR primers. PCR was performed on QuantStudio 5 Real-Time *PCR* System. Human cell engraftment percentage was calculated on the quantity of human gDNA within 100ng rat tissue per section.

### Lung Injury Score (LIS)

Hematoxylin and Eosin (H&E) staining was performed by the Molecular Pathology Core (MP/MPSR, Columbia University Medical Center) using a standard protocol. The slides were scanned using a bright field whole slide scanning system (Leica AT2 digital scanning system) with a pixel size 0.25µm. We then developed and applied a custom script to select random and non-overlapping regions of interest from whole slide image scans of lung sections. A section from the targeted left lobe and a section from the contralateral right lobe was used for the analysis. The script was used to obtained 30 regions of interest (ROI) per sample. Each ROI was then blindly evaluated for LIS according to guidelines of the 2011 ATS Workshop Report on Features and Measurements of Experimental Acute Lung Injury in Animals^38^ by a pulmonary pathologist (AS).

### Statistical analysis

All statistical tests are reported in the legends of the respective figures. All the statistical tests were performed using Prism v10 (GraphPad). A value of *p* < 0.05 was considered statistically significant.

## Author contributions

HYL, CP, HWS, and NVD designed the study, with advice from GVN and YWC. The cells are developed in the lab of HWS, the rat model in the lab of NVD. HYL performed all culture and in vitro analysis of DLEPs and analyzed rats by qPCR quantification. YWC first grew the DLEPs from lung organoids. CP performed all the animal experiments and relative histological analysis. MP analyzed scRNAseq data. SL contributed to the IF stainings of the rat lung sections. JW contributed to designing and optimizing the de-epithelialization protocol with CP and NVD. AS analyzed the LIS together with CP. SF contributed to troubleshooting the quantification of cell engraftment. JWM oversaw confocal imaging. HYL, CP, HWS, and NVD interpreted and analyzed the data. HWS and NVD co-wrote the manuscript with support of HYL, CP, AS, GVN and YWC. All the authors read and approved the final manuscript.

## Acknowledgments

The authors thank the following collaborators and supporters: Institute of Comparative Medicine veterinary staff including Dr. Rivka L. Shoulson and Willie V. Copeland for supporting the animal studies; Confocal and Specialized Microscopy Shared Resource of the Herbert Irving Comprehensive Cancer Center at Columbia University, funded in part through the NIH/NCI Cancer Center Support Grant P30CA013696 for the coding applied to lung sections for LIS; The Imaging Core of the Columbia Center for Human Development/Center for Stem cell Therapies, supported by NIH S10 OD032447 (HWS), the Columbia Center for Translational Immunology Flow Cytometry Core, supported in part by NIH S10OD020056; Afsar Barlas and the Molecular Cytology Core Facility at Memorial Sloan Kettering Cancer Research Center for performing the immunohistochemistry; Columbia Center for Translational Immunology including Adam D. Griesemer and Benjamin Piegari for tacrolimus quantification in immunosuppressed rats.

## Funding support

This work was supported by grants NIH HL120046 (HWS and GVN), NIH 1U01HL134760 (HWS and GVN), NIH S10 OD032447 (HWS), 1R01HL142727 (HWS), the Parker B. Francis Fellowship (YWC), the Thomas R. Kully IPF Research Fund (HWS), Driscoll Children’s Fund Scholar (NVD), Louis V. Gerstner Jr. Scholarship Fund (NVD), and by the Stony-Herbert Wold Fund (NVD).

## Data Availability

scRNAseq data are available in the GEO database (GEO Submission (GSE240574), access code: itsfowcmzzuxhyr).

## Competing interests

HWS, NVD, YWC and CP are inventors on pending patent applications related to this work.

## Extended data

**Extended Data Figure. 1.**
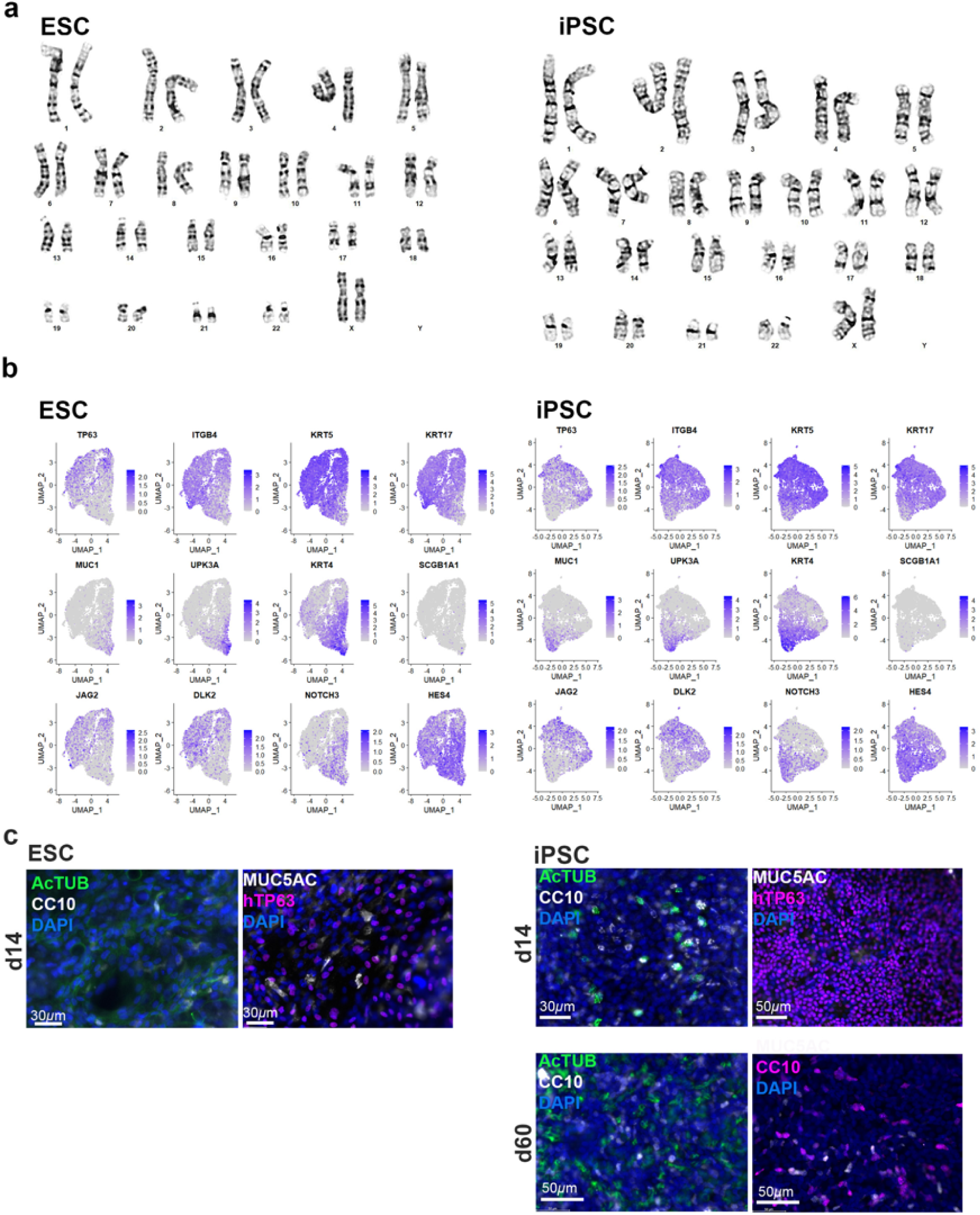
Characterization of secretory-like and Bc-like cells. **a**, Norm al karyotype (46, XX) of ESC-derive d and iPSC-derive d cells. **b**, UMA P feature plots for indicated markers in uninte grated analysis of ESC-derived and iPSC-derived cells. **c**, Expression of airway markers of ESC-derived cells on day 14 and iPSC-derived cells on day 14 and 60 in ALI culture conditions. **d**, Flow Strategy of gating single cells and subpopulation co-expressing EpCAM^+^ and MUC1^+^.

**Extended Data Figure. 2.**
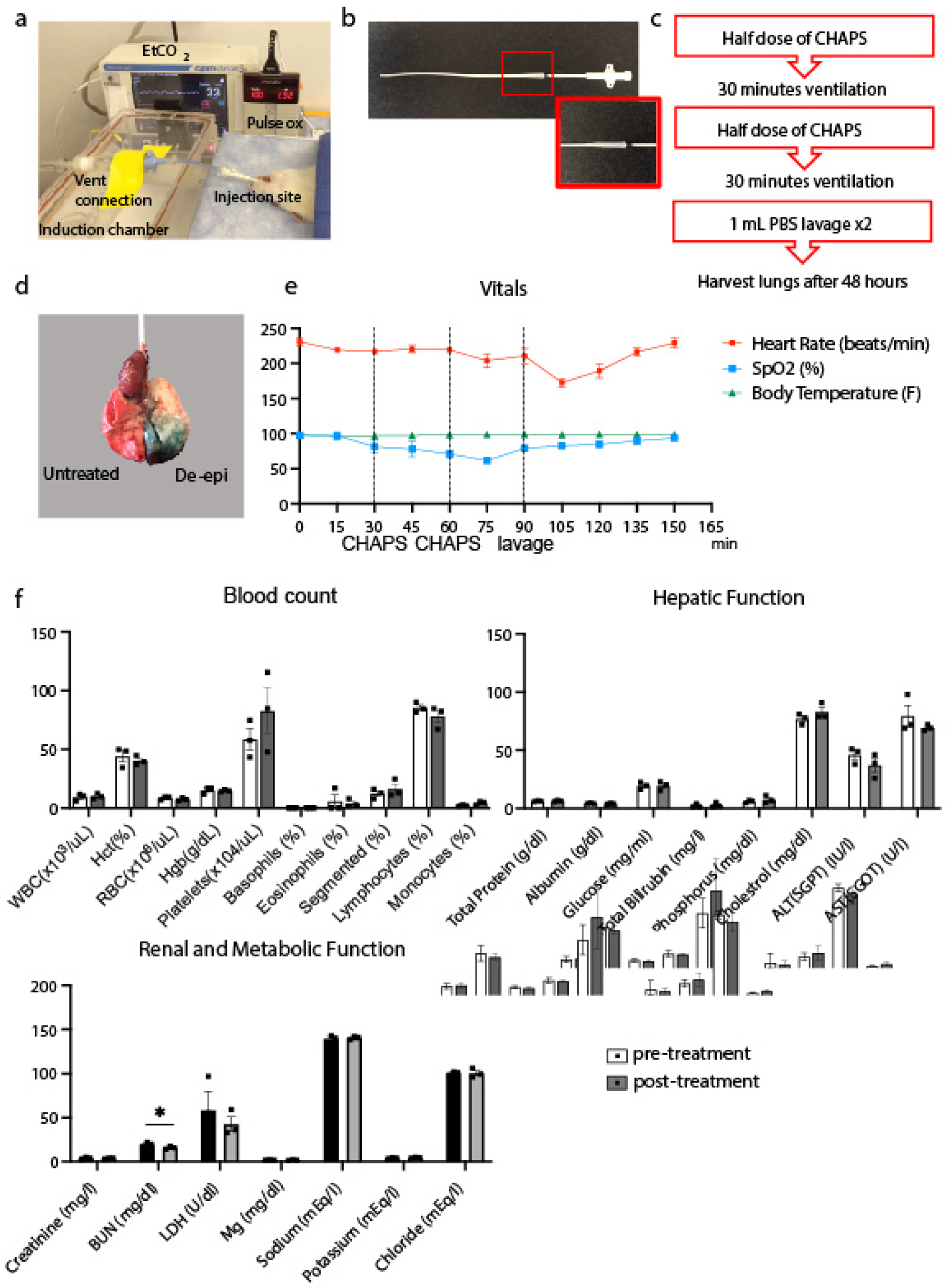
*In vivo* de-epithelialization. **a,** Experimental set up. **b,** Modified cannula for de-epithelialization. The red box highlights the tracheal plug. **c,** Schematic representation of the de-epithelialization procedure. **d,** Explanted lungs after intratracheal injection of a mixture of food blue dye with de-epithelialization solution. **e.** Heart rate (HR), oxygen saturation (SpO_2_), and rectal temperature (Temp) during the procedure. Mean±SEM (n=3). **f**. Peripheral blood analysis before (Pre-) and 48 hours after (Post-) treatment. Mean±SEM (n=3). Two-tailed paired Student’s t-tests (*, p = 0.02).

**Extended Data Figure. 3.**
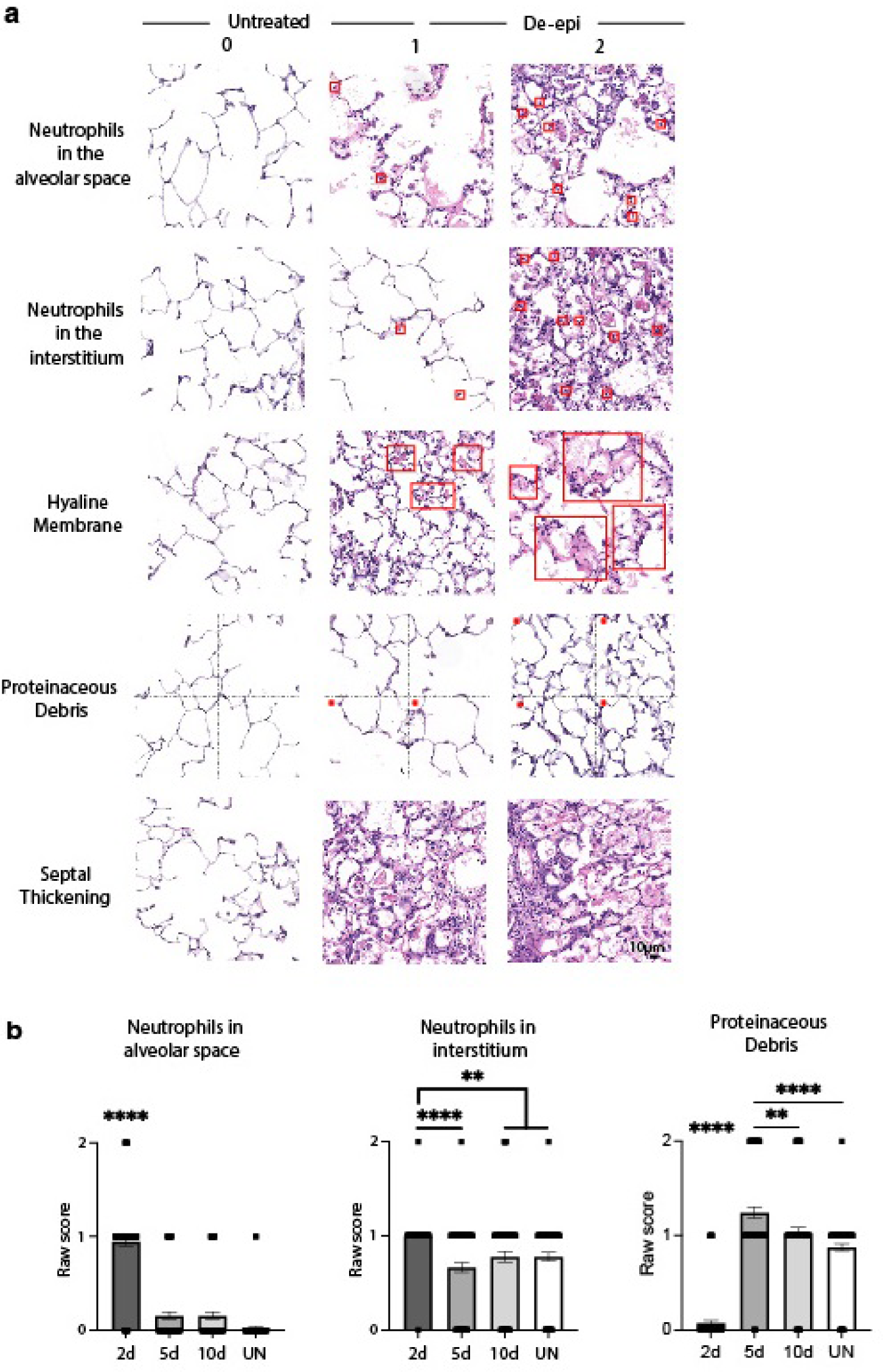
LIS individual features after de-epithelialization injury. **a.** Examples of scoring evaluation from each LIS feature. Red squares in ROI highlight neutrophils in the first two rows, and hyaline membranes in the third row. Dotted black lines divide the ROI in quadrants for the evaluation of the presence of proteinaceous debris in forth row. **b.** Scores for individual features of the LIS. Mean±SEM, paired t-test, n=3 animals per condition, 60 ROI (30 from left lung, 30 from right lung) per animal; **, p=0.0037-0.0081; ****, p<0.0001.

**Extended Data Figure. 4.**
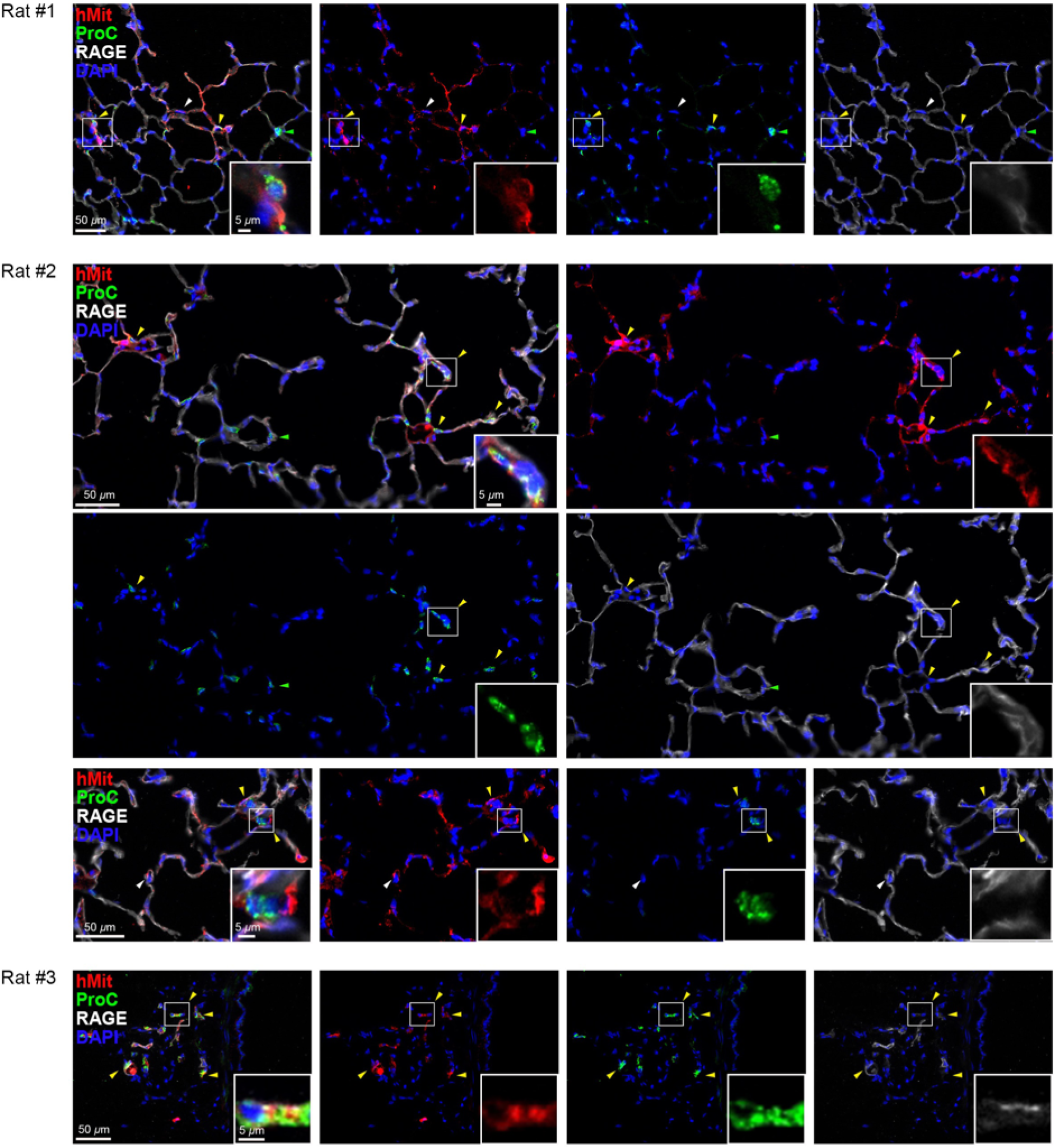
Expression of Pro-SPC in engrafted DLEPs in rats conditioned with de-epithelialization on day 10. Representative areas showing engraftment of human cells based on hMit staining in different rats conditioned with de-epithelialization (White arrowheads: examples of human cells co-expressing hMit and RAGE. Yellow Arrowheads: human cells co-expressing hMit and ProSPC. Green arrowheads: native rat ProSPC^+^ cells). Single channels and a representative higher magnification area of engrafted cells are highlighted in the white squares.

**Extended Data Figure. 5.**
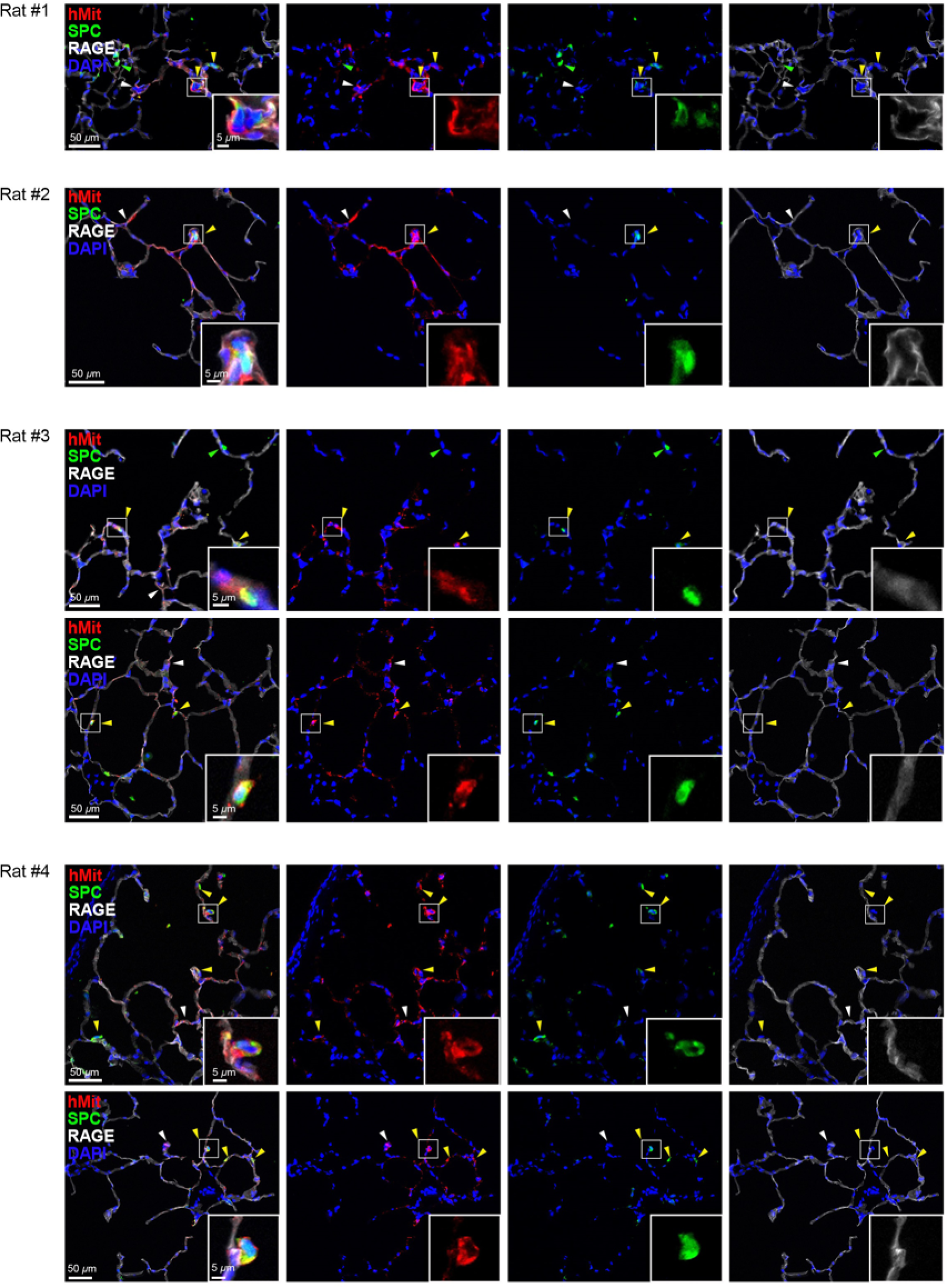
Expression of SPC in engrafted DLEPs in rats conditioned with de-epithelialization on day 10. Representative areas showing engraftment of human cells based on hMit staining in different rats conditioned with de-epithelialization (White arrowheads: examples of human cells co-expressing hMit and RAGE. Yellow Arrowheads: human cells co-expressing hMit and ProSPC. Green arrowheads: native rat ProSPC^+^ cells). Single channels and a representative higher magnification area of engrafted cells are highlighted in the white squares.

**Extended Data Figure. 6.**
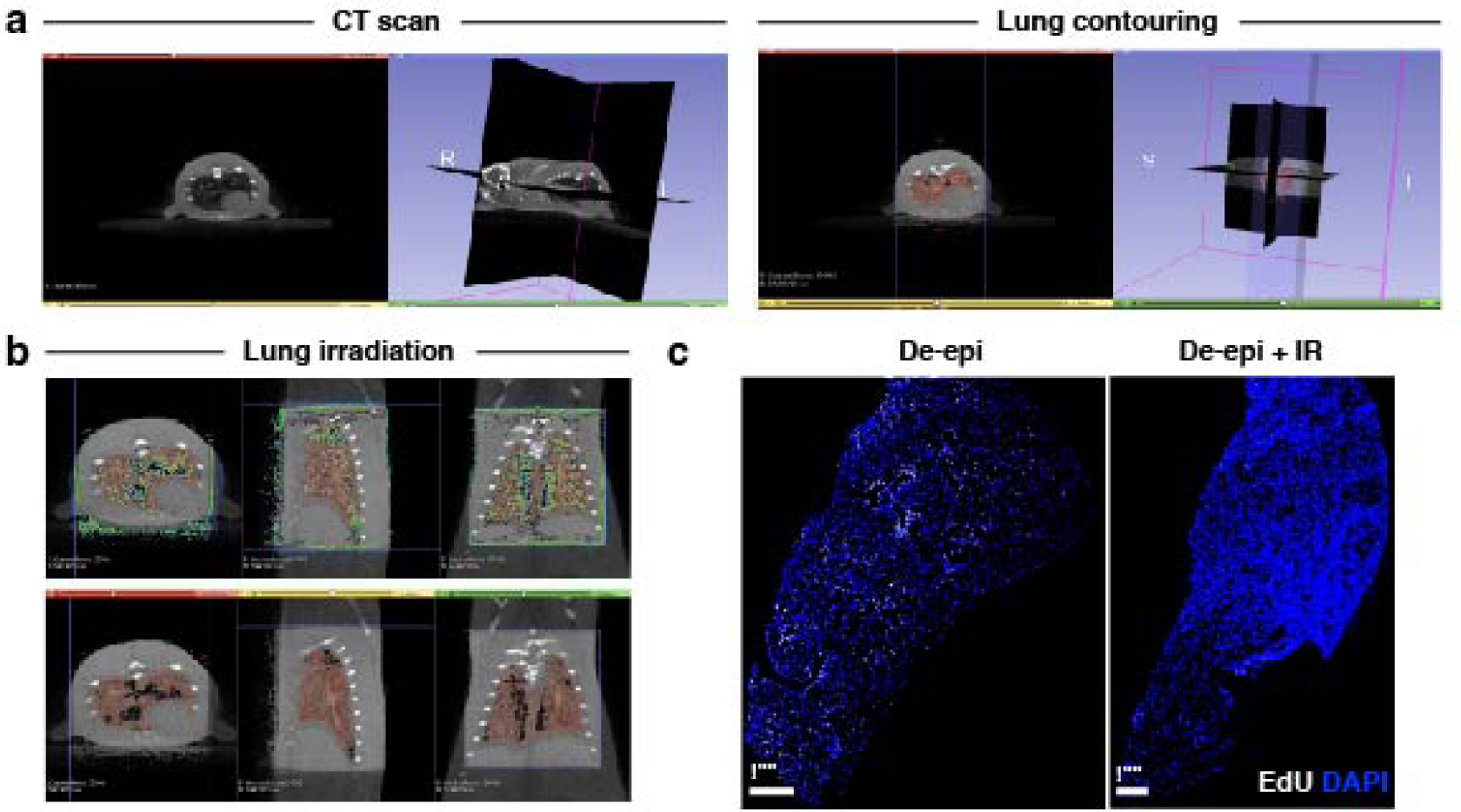
Rat lung irradiation. **a.** CT scan and lung contouring. **b.** Lung irradiation field (contoured regions). **c.** EdU incorporation after de-epithelialization with and without irradiation.

**Extended Data Figure. 7.**
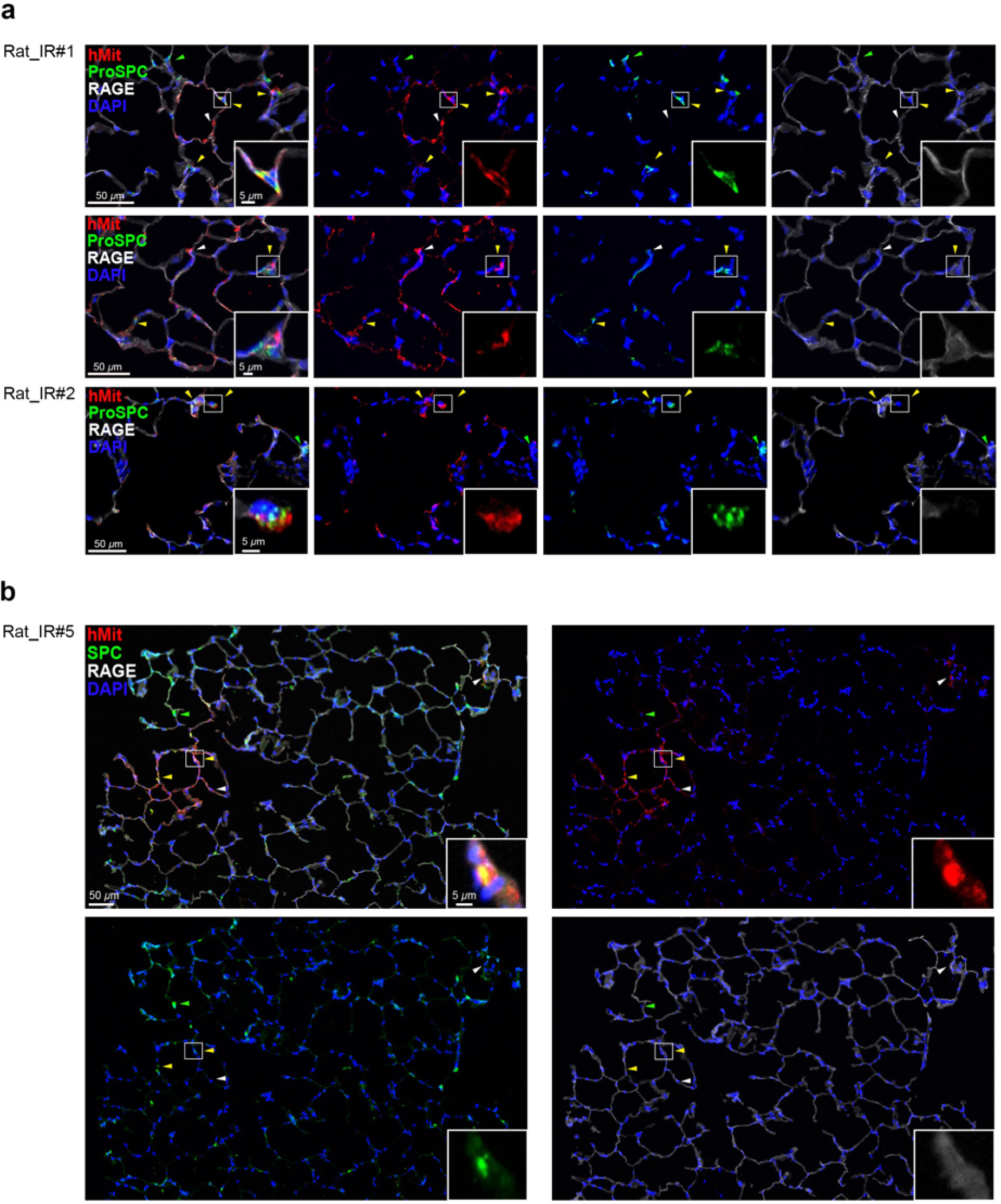
Expression of Pro-SPC and SPC in engrafted DLEPs in rats conditioned with de-epithelialization and irradiation on day 10. Representative areas showing engraftment of human cells based on hMit staining in different rats conditioned with de-epithelialization and (White arrowheads: examples of human cells co-expressing hMit and RAGE. Yellow Arrowheads: human cells co-expressing hMit and ProSPC (**a**) or SPC (**b**). Green arrowheads: native rat ProSPC^+^ (**a**) or SPC^+^ cells(**b**)). Single channels and a representative higher magnification area of engrafted cells are highlighted in the white square.

**Extended Data Figure. 8.**
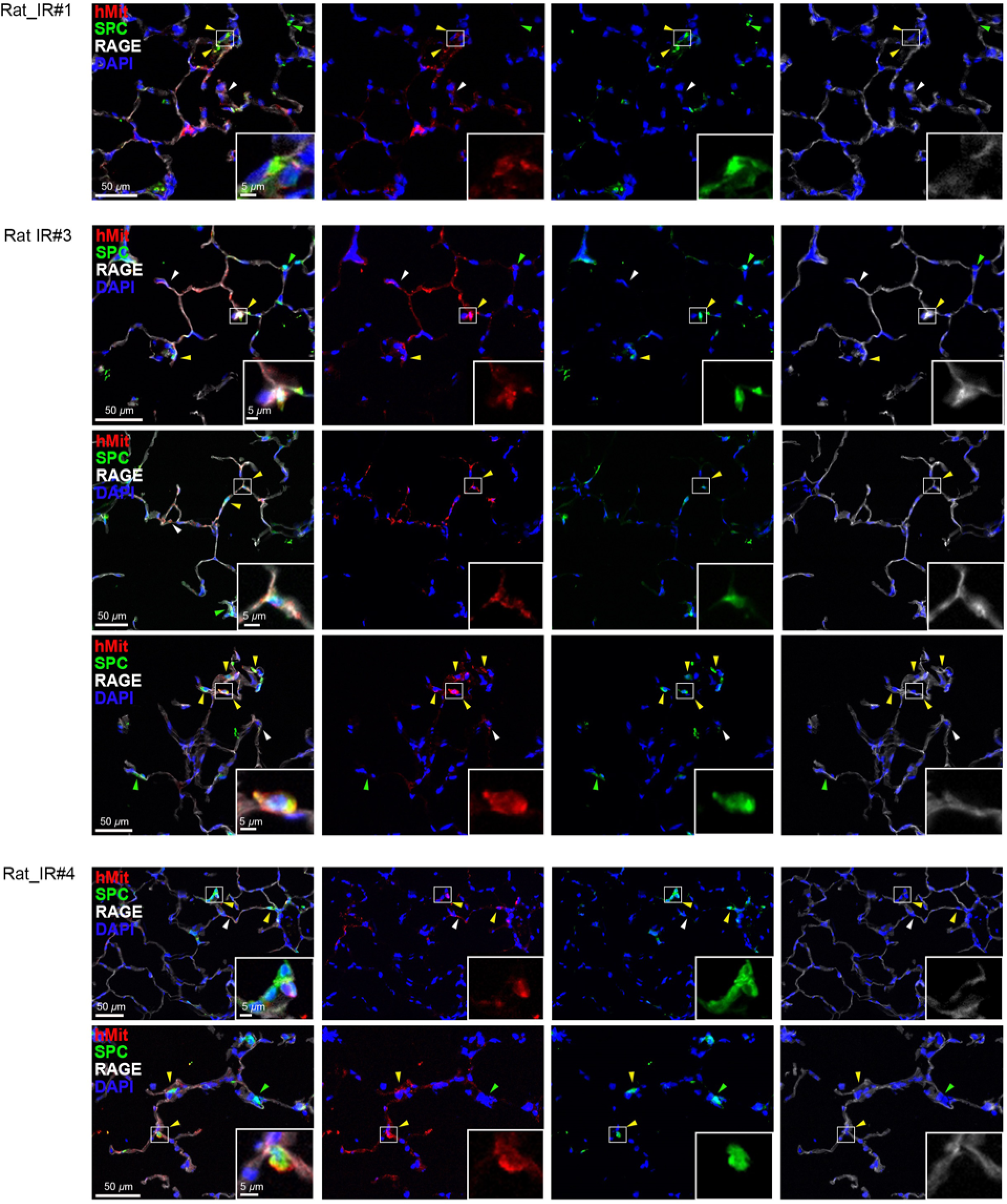
Expression of SPC in engrafted DLEPs in rats conditioned with de-epithelialization and irradiation on day 10. Representative areas showing engraftment of human cells based on hMit staining in different rats conditioned with de-epithelialization and irradiation (White arrowheads: examples of human cells co-expressing hMit and RAGE. Yellow Arrowheads: human cells co-expressing hMit and SPC. Green arrowheads: native rat SPC^+^ cells). Single channels and a representative higher magnification area of engrafted cells are highlighted in the white square.

**Extended Data Figure. 9.**
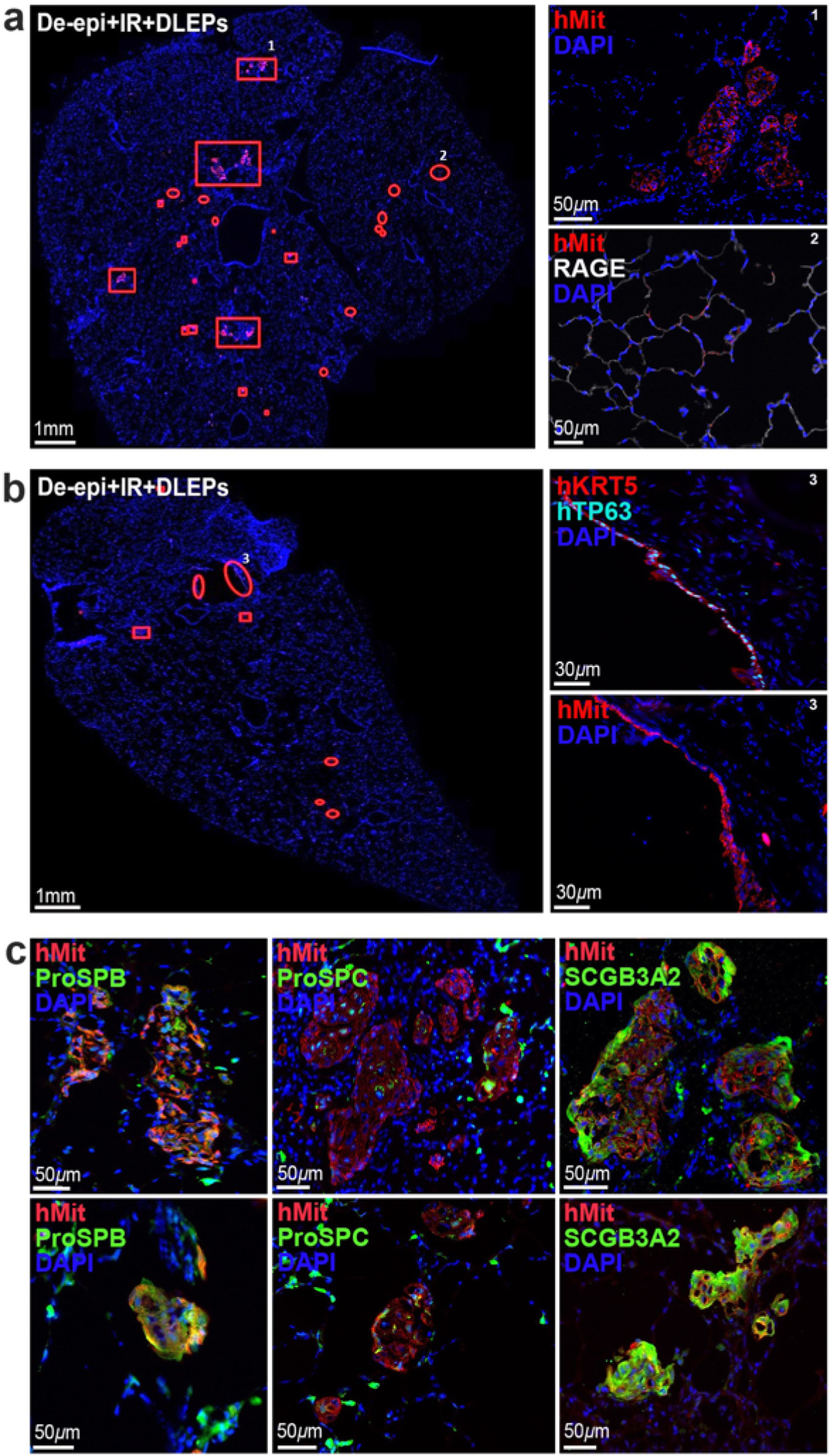
Engraftment of DLEPs after de-epithelialization and irradiation on day 10. **a.** Annotations indicating location of human cells based on hMit staining. Rectangles: engraftment in KRT5-pods; circles: engraftment in alveolar network. **b.** Annotation indicating location of human cells based on hMit staining. Rectangles: engraftment in pods; ellipses: engraftment in airways. **c.** Expression of hMit, ProSPB, ProSPC and SCG3A2 in engrafted KRT5-pods.

**Extended Data Figure. 10.**
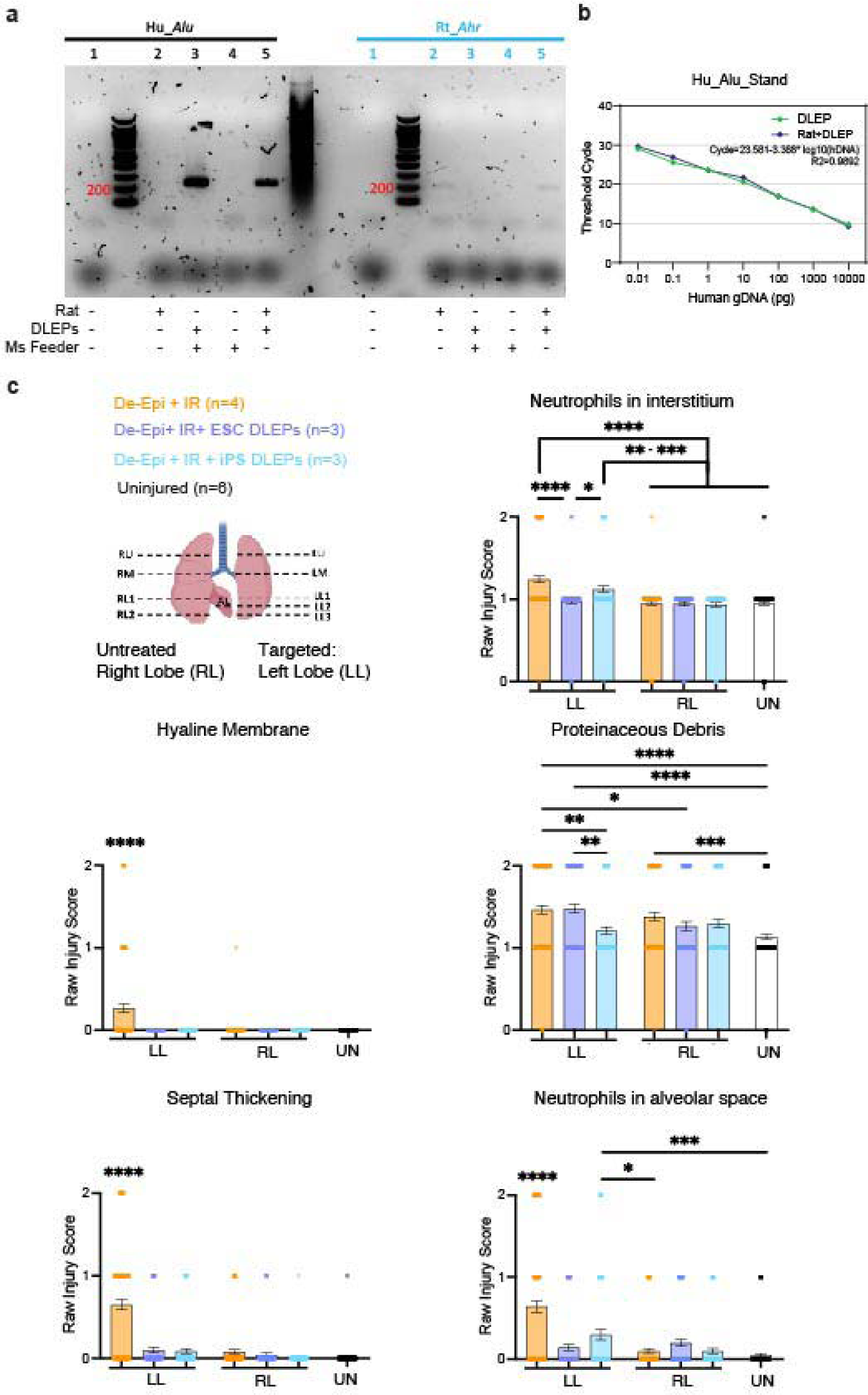
Quantification of DLEP engraftment and lung injury repair. **a.** Gel electrophoresis of PCR products obtained after amplification of rat lung tissue, DLEPs, and mouse feeder cells with AluY8b and rat Ahr primer sets. **b.** Standard curves for AluY8b human primers in DLEPs only and DLEPs mixed with 100ng rat lung gDNA. **c**. Scores for individual features of the LIS. Mean±SEM (IR: irradiation, UN: normal control lung, 800 blindly evaluated fields in total, 30 ROI per tile scan, one-way ANOVA; *, p=0.0301-0.0466; **, p=0.0014-0.0040; ***, p=0.0002-0.0009]. ****, p<0.0001].

## Notes

### Competing Interest Statement

HWS, YWC, CP and NVD have filed patent applications relevant to the work presented.

